# Metal-driven anaerobic oxidation of methane as an important methane sink in methanic cold seep sediments

**DOI:** 10.1101/2022.12.21.518016

**Authors:** Xi Xiao, Min Luo, Chuwen Zhang, Tingting Zhang, Xiuran Yin, Xuemin Wu, Jing Zhao, Jun Tao, Zongheng Chen, Qianyong Liang, Xiyang Dong

## Abstract

Anaerobic oxidation of methane (AOM) coupled with reduction of metal oxides is supposed to be a globally important bioprocess in marine sediments. However, the responsible microorganisms and their contributions to methane budget are not clear in deep sea cold seep sediments. Here, we combined geochemistry, muti-omics and numerical modeling to study metal-dependent AOM in methanic cold seep sediments in the northern continental slope of the South China Sea. Geochemical data based on methane concentrations, carbon stable isotope, solid-phase sediment analysis and pore water measurements indicate the occurrence of anaerobic methane oxidation coupled to metal oxides reduction in the methanic zone. The 16S rRNA gene amplicons and transcripts, along with metagenomic and metatranscriptomic data suggest that diverse ANME groups actively mediated methane oxidation in the methanic zone either independently or in syntrophy with e.g. ETH-SRB1 as potential metal reducers. Modeling results suggest that the estimated rates of methane consumption via Fe-AOM and Mn-AOM were both 0.3 μmol cm^-2^ yr^-1^, which account for ∼3% of total CH_4_ removal in sediments. Overall, our results highlight metal-driven anaerobic oxidation of methane as an important methane sink in methanic cold seep sediments.

## Introduction

Marine cold seeps are not only an indicator of gas hydrate reservoirs, but also an important methane source to the oceans, which has a significant impact on the global carbon cycle and climate change^1^. It is estimated that 0.02 Gt methane is consumed annually in the sediment with an additional 0.02 Gt methane releasing annually into the overlying ocean by seafloor cold seeps^2^. Thus, the production and consumption of methane is a key component of the carbon cycle for cold seeps^3^. Anaerobic oxidation of methane (AOM) driven by microbial communities, plays a key role in decreasing methane emissions to the atmosphere^4^. Anaerobic methanotrophic archaea (ANME) mediate this process through the coupling of methane oxidation to the reduction of nitrite, nitrate, manganese/iron oxides, and sulfate^5-8^. Sulfate-driven AOM (S-AOM) by assemblages of ANME and sulfate-reducing bacteria (SRB)^6, 9^ is regarded as the major process for methane sink within cold seep sediments, reaching highest activities within the sulfate-methane transition zone (SMTZ)^10^.

Early studies show that methane oxidation possibly coupled with metal oxidation can still occur at a considerable rate in the methanic zone below the SMTZ when sulfate has been completely depleted or at very low levels^11^. A range of efforts have been undertaken to demonstrate the occurrence of metal oxides-driven AOM (Metal-AOM, including high-valence iron/manganese oxides)^5, 12, 13^. This microbial process is mediated by ANME through the reverse methanogenesis pathway, typically in syntrophy with dissimilatory iron/manganese reducing bacteria^5, 14^. Investigations of enrichment cultures have also revealed that ANME-2a, ANME-2c, and ANME-2d can perform AOM coupled to the extracellular dissimilatory reduction of iron and manganese oxides independently using e.g. a unique set of multiheme cytochromes (MHCs)^12, 13, 15-17^.

The activity rates of Fe-AOM are efficiently estimated by incubation experiments or geochemical modeling, but rarely for Mn-AOM (**Table 1**)^18-25^. Microbial culture experiments from the Eel River Basin seep have found that manganese oxides can drive AOM as electron acceptors more efficiently than ferrihydrite^5^. However, microorganisms involved in the coupling between AOM and metal reduction in marine environments are still largely unknown, especially for manganese reduction^26^. The contribution of Mn-AOM for CH_4_ removal in situ marine environments is still fully identified, neither. Without in-depth understanding of the role of Metal-AOM in the biogeochemical cycle, the contribution of metal reduction to the global carbon cycle, especially the methane sink, is likely to be undervalued^27^.

**Table 1.** Summary of the estimated rates of S-AOM, Fe-AOM and Mn-AOM in sediments from various freshwater and marine environments.

| Ecosystem | Environment | In situ concentration (μmol/L) |  | Model-derived rates (μmol CH<sub>4</sub> cm<sup>-3</sup> yr<sup>-1</sup>) |  |  | Depth-integrated rates (μmol CH<sub>4</sub> cm<sup>-2</sup> yr<sup>-1</sup>) |  |  | The fraction in total CH<sub>4</sub> oxidation |  |  | Method | References |
| --- | --- | --- | --- | --- | --- | --- | --- | --- | --- | --- | --- | --- | --- | --- |
|  |  | Fe<sup>2+</sup> | Mn<sup>2+</sup> | S-AOM | Fe-AOM | Mn-AOM | S-AOM | Fe-AOM | Mn-AOM | S-AOM | Fe-AOM | Mn-AOM |  |  |
| Marine | Haima cold seep | 148 | 2289 | 0.66 | 0.02 | N.A. | 20.05 | 0.31 | 0.32 | 97% | 1.5% | 1.5% | Modeling | This study |
|  | Hikurangi margin | 184 | N.A. | 0.48 | 0.0005 | N.A. | 3 | 0.4 | N.A. | 88% | 12% | N.A. | Modeling | Luo et al., 2020 |
|  | Black Sea | 800 | 23 | 0.07 | 1.46E-05 | N.A. | 5.9 | 0.04 | N.A. | 99% | 0.70% | N.A. | Modeling | Egger et al., 2016a |
|  | Baltic Sea | 600 | N.A. | 0.27 | 0.0011 | N.A. | 8.8 | 2.5 | N.A. | 78% | 22% | N.A. | Modeling | Egger et al., 2017 |
|  | Bothnian Sea | 1830 | N.A. | N.A. | N.A. | N.A. | 78.7 | 8 | N.A. | 90% | 9% | N.A. | Modeling | Rooze et al., 2016 |
|  |  |  |  | 6.94E-04 | 1.32 | N.A. | 58.38 | 1.63 | N.A. | 97% | 3% | N.A. | <sup>13</sup>CH<sub>4</sub>/Modeling | Egger et al., 2015 |
|  | Jiaolong cold seep | 27 | N.A. | 328.57 | 13.87 | N.A. | N.A. | N.A. | N.A. | N.A. | N.A. | N.A. | <sup>14</sup>CH<sub>4</sub> | Li et al., 2020 |
|  | North Sea | 380 | 40 | 2.04 | 0.03 | N.A. | N.A. | N.A. | N.A. | 98% | 2% | N.A. | <sup>14</sup>CH<sub>4</sub> | Aromokeye et al., 2020 |
|  | Eel River Basin seep | N.A. | N.A. | 52 | 6 | 14 | N.A. | N.A. | N.A. | N.A. | N.A. | N.A. | <sup>13</sup>CH<sub>4</sub> | Beal et al., 2009 |
| Freshwater | Lake Kinneret | 70 | N.A. | N.A. | 1.26 | N.A. | N.A. | N.A. | N.A. | N.A. | N.A. | N.A. | <sup>13</sup>CH<sub>4</sub> | Sivan et al., 2011 |
|  | Dover Bluff salt marsh | 30 | 80 | 2.41 | 1.42 | 0.876 | N.A. | N.A. | N.A. | N.A. | N.A. | N.A. | <sup>14</sup>CH<sub>4</sub> | Segarra et al., 2013 |
|  | Hammersmith Creek River | 500 | 400 | 5.66 | 4.5 | 1.314 | N.A. | N.A. | N.A. | N.A. | N.A. | N.A. | <sup>14</sup>CH<sub>4</sub> |  |

Modeling: Geochemical modeling estimates; <sup>13</sup>CH<sub>4</sub>: <sup>13</sup>CH<sub>4</sub> incubations; <sup>14</sup>CH<sub>4</sub>: <sup>14</sup>CH<sub>4</sub> incubations; N.A., not available.

Haima cold seep was firstly discovered as an active cold seep in the Qiongdongnan basin on the northwest slope of the South China Sea by the dives of a remotely operated vehicle named as Haima in 2015 (**Figure S1a**)^28^. A large number of findings have since been emerging about the biogeochemistry of cold-seep carbonates, benthos and sediments in Haima cold seep^29-32^. Massive amounts of terrigenous metal oxides are supplied into the continental slope of the South China Sea from rivers^33^. Consequently, iron/manganese-containing minerals are part of the major components in the sediments of this region with high-flux methane seeps, rendering it be such a natural laboratory to investigate the role of metal oxides in methane cycle.

In this study, combining geochemical and microbial analyses of the Haima cold seep sediments, we aimed to (1) reveal the occurrence of Metal-AOM in the methanic zone; (2) identify microorganisms involved in the Metal-AOM and their key mechanisms; (3) estimate the contribution of removal of methane by Fe/Mn-AOM. Our findings provide insights into the coupling mechanism between iron/manganese reduction and AOM as well as the role of Metal-AOM in the biogeochemical cycle.

## Results and Discussions

### Geochemical data indicate anaerobic methane oxidation in the methanic zone

At a water depth of 1375 m, a 4.3-meter-long piston core was retrieved from the Haima cold seep in the South China Sea, where a gas chimney and bubble plumes were observed indicative of ongoing seepage activities (**Figure S1b-d**). Massive gas hydrates were also found at 347-352 cm below the seafloor (cmbsf) and 420 cmbsf (**Figure S1e-g**). Methane (CH_4_) was the dominant seeping hydrocarbon gas (0.13-919.57 μM), along with ethane (C_2_H_6_) being detected (0.07-3.96 μM) below 130-140 cmbsf (**Table S1**). Based on methane and sulfate profiles (**Figure 1a and Table S1**), our sediment core samples were categorized into three biogeochemical zones^34, 35^: sulfate reduction zone (surface-130 cmbsf), sulfate–methane transition zone (∼130-250 cmbsf), and methanic zone (250-430 cmbsf).

**Figure 1.**
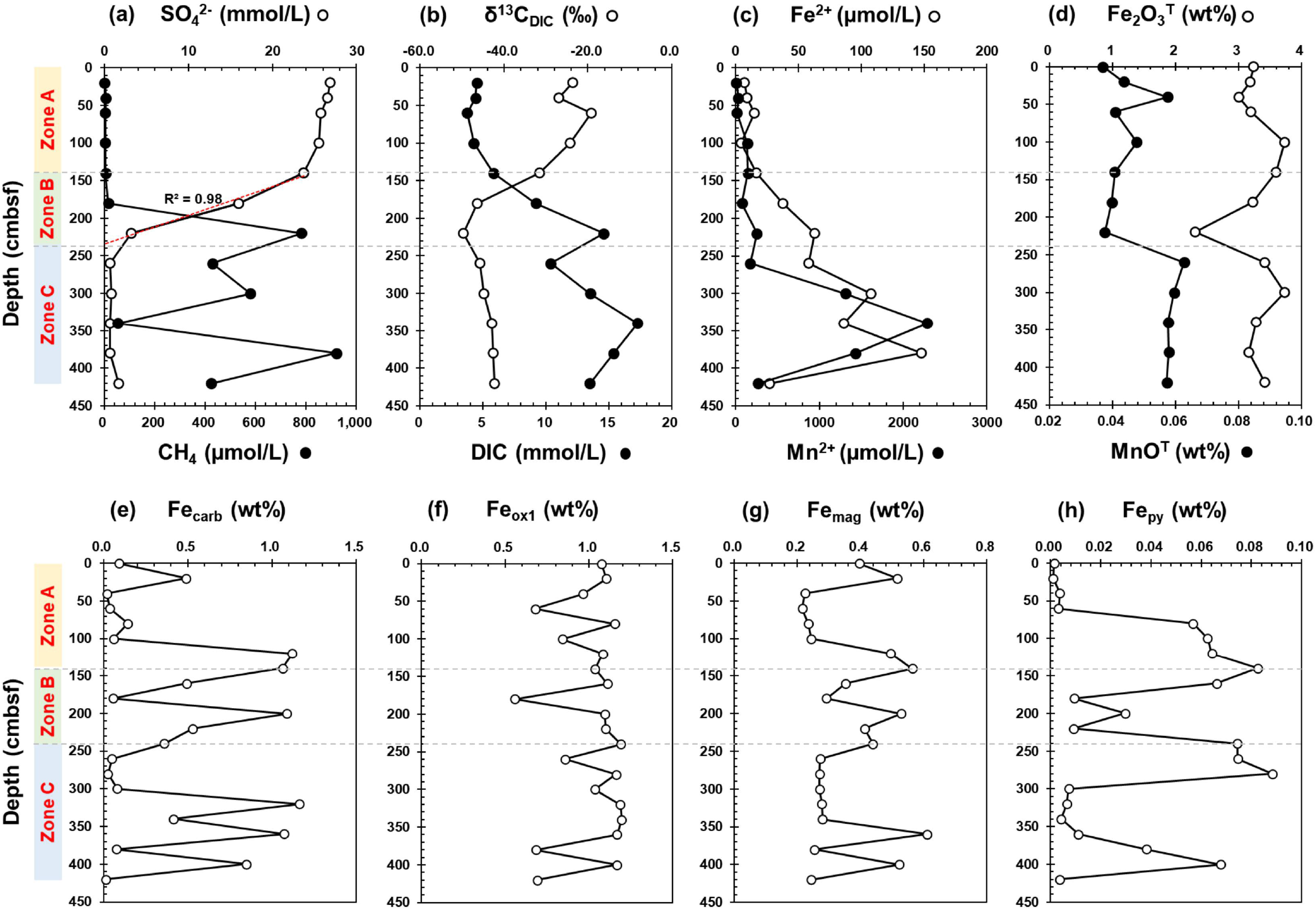
Geochemical profiles of the sediment core HM-S11 in the Haima seep. (a) Profiles of methane (CH_4_) and sulfate (SO_4_^2−^) contents in porewater; (b) Concentrations of dissolved inorganic carbon (DIC) and stable carbon isotope ratios δ^13^C_DIC_ in porewater; (c) Concentrations of dissolved Fe^2+^ and Mn^2+^ in porewater; (d) Contents of Fe_2_O_3_^T^ and MnO_2_^T^ in sediments; (e∼h) Sequential extraction of iron minerals in sediments. Fe_carb_ = carbonate-associated Fe; Fe_ox1_ = amorphous iron (oxyhydr)oxides; Fe_mag_ = magnetite Fe; Fe_py_ = pyrite Fe. Zones A, B and C are suggested as the sulfate zone, the sulfate-methane transition zone, and the methanic zone.

In the methanic zone, methane concentrations fluctuated between 53 μM and 920 μM. A notable decrease was observed from 781 μM at 210-220 cmbsf to 53 μM at 350 cmbsf, with the enrichment of the stable carbon isotope ratios (δ^13^C) in residual CH_4_ from −68.77‰ to −64.33‰ (**Table S1**), pointing to occurrence of biological unitization of methane^25^. In accordance with this, the dissolved inorganic carbon (DIC) values increased from 250-260 cmbsf (10.38 mM) to 330-340 cmbsf (17.27 mM; **Figure 1b**), as also evidenced by the increase in total alkalinity^36^ from 16.01 mM to 28.94 mM (**Table S1**). The measured δ^13^C_DIC_ values (<−42.30‰; **Figure 1b**) were much more ^13^C-depleted than that of typical marine organic matters (∼−20‰) in this sea area^37^. This confirms that DIC increase was caused by microbial methane oxidation rather than microbial degradation of other organic matters. Additionally, low concentration profiles of phosphate (PO_4_^3−^) and ammonium (NH_4_^+^), lower than 41.65 μM and 56.72 μM respectively (**Table S1**) also support that organic matter degradation was not the main reason for increased DIC concentrations in these sediment samples^38^.

### Diverse ANME actively mediated methane oxidation in the methanic zone

Anaerobic methanotrophs are assigned to three distinct clades (ANME-1 with subgroups a and b, ANME-2 with subgroups a, b, c and d, and ANME-3) within the phylum Halobacteriota^39^. To identify potential ANME clades in the methanic zone, we performed DNA and RNA sequencing of sediment samples. Taxonomy classifications of archaeal 16S rRNA gene amplicons indicate that ANME accounted for 69∼87% of the whole archaeal community in the methanic zone (**Figure 2a and Table S2**). These ANME populations are phylogenetically diverse, including ANME-1a (up to 80% at 320-330 cmbsf), ANME-1b (up to 24% at 420-430 cmbsf), ANME-2c (up to 16% at 280-290 cmbsf) and ANME-3 (up to 68% at 360-370 cmbsf). Relative abundances of 16S rRNA transcripts suggest that ANME-1a (up to 54% at 320-330 cmbsf), ANME-3 (up to 84% at 360-370 cmbsf), ANME-2c (up to 63% at 280-290 cmbsf), and ANME-1b (up to 19% at 400-410 cmbsf) were the dominant and active players for AOM occurring in the methanic zone (**Figure 2b and Table S2**).

**Figure 2.**
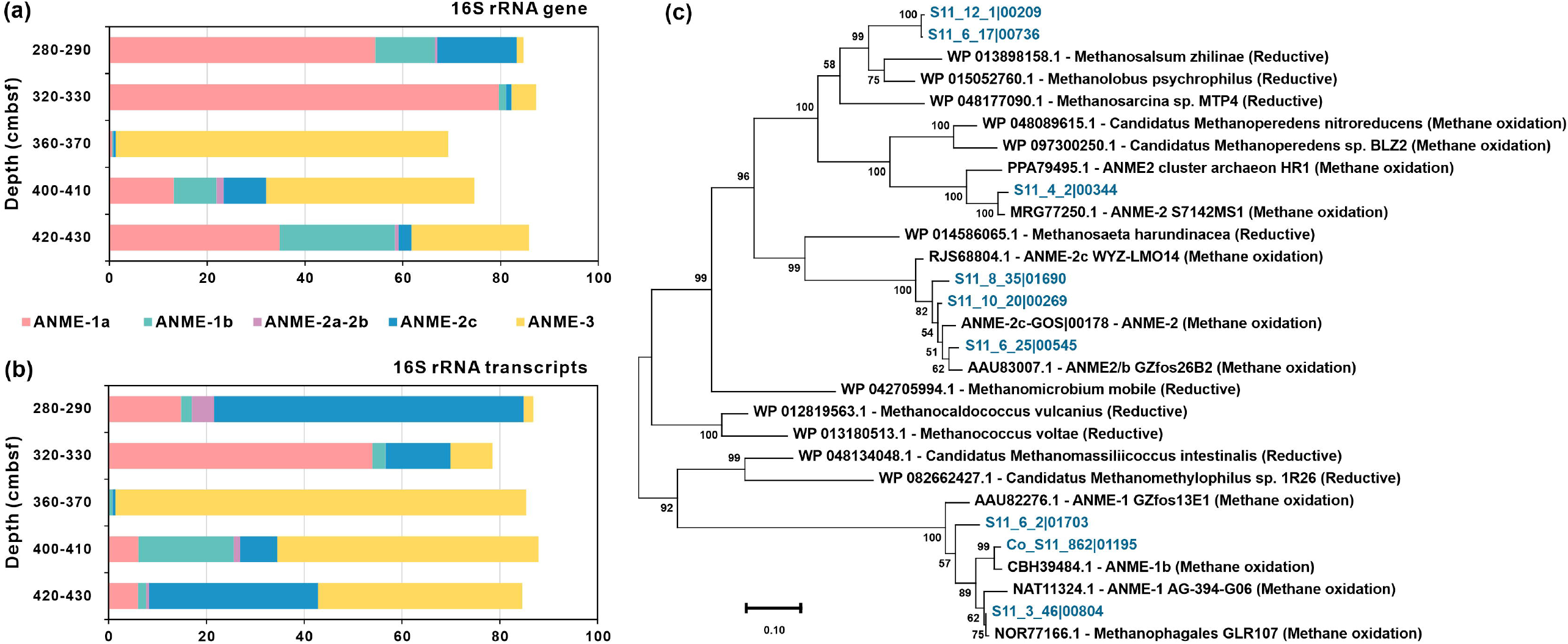
Relative abundances and phylogenetic tree of ANMEs. Relative abundances of ANME based on (a) 16S rRNA genes and (b) 16S rRNA transcripts in the methanic zone. (c) Maximum-likelihood phylogenetic tree of *mcrA* sequences.

Metagenomic assembly and binning yielded 17 metagenome-assembled genomes (MAGs) taxonomically affiliated with methane-metabolizing lineages that either produce or consume methane (**Table S3**). Among them, nine harbor sequences encoding the catalytic subunit of methyl-coenzyme M reductases (McrA)^40^ involved in methyl reduction during methane oxidation (**Figure 2c and Table S4**). They belong to clades of ANME-1 (n=3) and ANME-2 (n=4). Additionally, S11_12_1 and S11_6_17 belonging to *Methanosarcinaceae* were predicted to have the capability to perform the methanogenesis pathway. MAGs for ANME-3 lineages were not recovered despite its highly relative abundance based on 16S rRNA genes and transcripts (**Table S2**). Based on read mapping (**Table S3**), species represented by ANME-1 (i.e., Ca. *Methanophagales*^41^) S11_3_46 (0.4-7.7 % of the whole microbial community) and S11_6_2 (0.4-7.3%) were observed to be the most abundant in the methanic zone, followed by S11_10_20 (0.1-1.2%) and S11_8_35 (0.1-0.6%) from ANME-2c (Ca. *Methanogaster*^39, 40^). Metatranscriptomic analyses (**Table S5**) showed that three ANME-1 genomes (Co_S11_862, S11_3_46 and S11_6_2) highly expressed *mcrA* genes in methanic zone (up to 4382 TPM at 400-410 cmbsf). The transcripts of *mcrB* genes from ANME-1 (Co_S11_862) and ANME 2c (S11_12_8) genomes also had high expression levels (up to 2378 TPM at 400-410 cmbsf). These results further suggest that ANME populations were actively responsible for the observed anaerobic methane oxidation in the methanic zone.

### Methane oxidation is coupled to metal oxides reduction in the methanic zone

In the methanogenic zone, dissolved ferrous iron (Fe^2+^) and manganese (Mn^2+^) concentrations in pore water were found to reach up to 148 μM at 370-380 cmbsf and as high as 2289 μM at 330-340 cmbsf, respectively (**Figure 1c and Table S1**). The Spearman correlations (**Figure 3a**) results further show that Fe^2+^ and Mn^2+^ concentrations have a strong positive covariance with CH_4_ (ρ = 0.874 and 0.699, respectively), DIC (ρ = 0.891 and 0.818, respectively); Fe^2+^ has a strong negative relationship with δ^13^C_DIC_ (ρ = −0.655; *p* < 0.05). These data indicate that the high amounts of dissolved Fe^2+^ and Mn^2+^ are associated with the fluctuation of methane concentrations in methanic zone^15, 42^. Correspondingly, the solid-phase sediment analysis revealed richer Fe_2_O_3_^T^ (3.17–3.74%) and MnO_2_^T^ (0.06%) in the methanic zone than those of the SMTZ (Fe_2_O_3_^T^: 2.31-3.60%; MnO_2_^T^: 0.04%) (**Figure 1d and Table S6**). The reactive iron minerals, including carbonate-associated iron (Fe_carb_; up to 1.16%), amorphous iron (oxyhydr)oxides (Fe_ox1_; up to 1.19%) and magnetite iron (Fe_mag_; up to 0.61%), were detected with the higher contents in the methanic zone (**Figure 1e-1g and Table S7**). Therefore, these data implied sufficient supplies of reactive Fe-oxides and the occurrence of Fe authigenic minerals as the products of iron reduction^43^. Similar to that of iron, total manganese (MnO_2_^T^) is also elevated from 0.04% in the SMTZ to 0.06% in the methanogenic zone (**Figure 1d and Table S6**). Given the elevated MnO_2_^T^ and extremely high dissolved Mn^2+^ (up to 2289 μM), the contribution of manganese reduction to AOM cannot be ignored in this seep. Overall, porewater and solid-phase profiles support metal-driven methane oxidation in methanic sediments.

**Figure 3.**
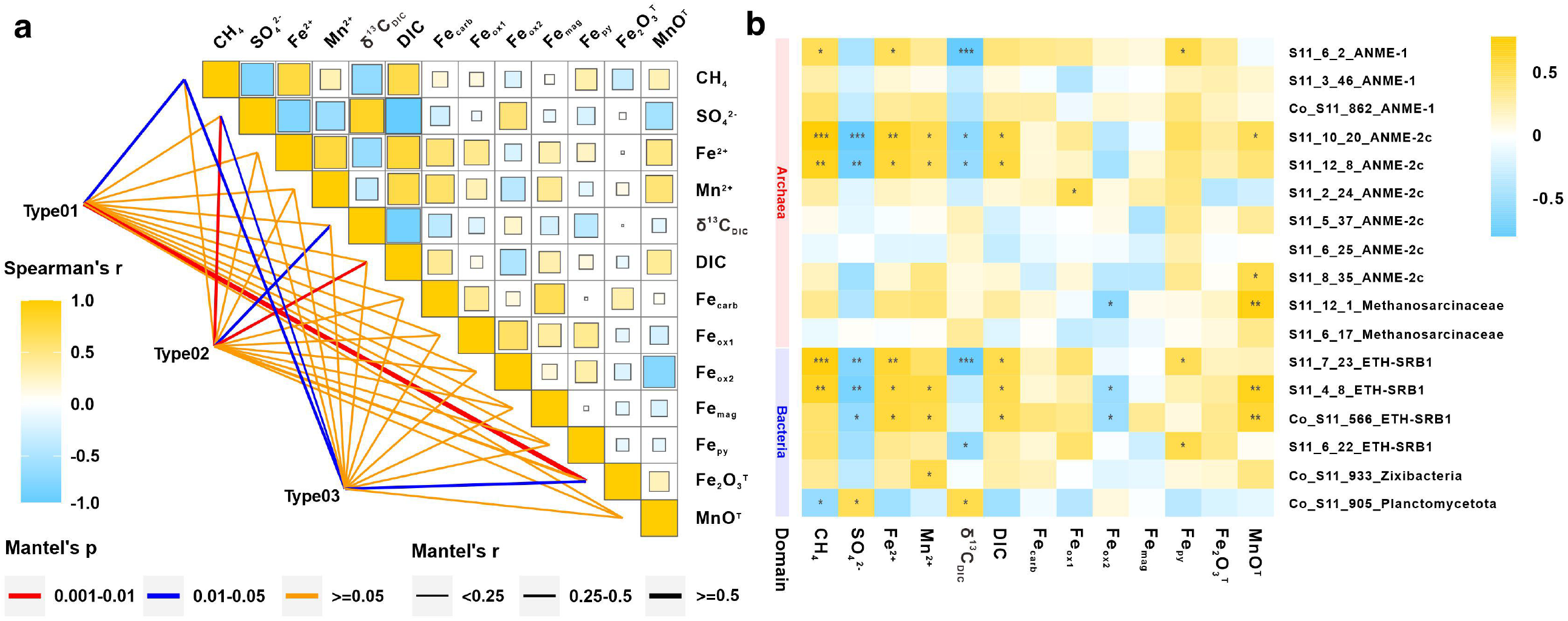
Spearman correlations of sediments in the core HM-S11. (a) Spearman correlation coefficients between depth-wise distribution of geochemical parameters; (b) Correlation between geochemical parameters and abundances of MAGs belong to ANMEs and metal reduction bacteria. The stars symbolize p values of correlation. *** means p < 0.001; ** means p < 0.01; *means p < 0.05.

### Potential microorganisms involved in dissimilatory metal reduction

For in-depth understanding of Fe(III)/Mn(IV)-dependent AOM in Haima cold seep, it is critical to identify the indigenous microorganisms responsible for this process. Members in different ANME clades are suggested to mediate metal-driven AOM by extracellular electron transfer (EET) to Mn(IV)/Fe(III) (oxyhydr)oxides or metal-reducing partners. In iron-reducer *Geobacter sulfurreducens*, the process of EET is carried out via MHCs during metal reduction^44 26^. For ANME-2d from freshwater Fe-AOM enrichment, a set of MHCs for extracellular dissimilatory Fe(III) reduction were highly expressed^12, 45, 46^. Here, all analyzed ANME genomes were found to contain the genes encoding several c-type and periplasmic cytochromes (**Table S8**). Among all ANME genomes, three MAGs, S11_2_24 and S11_6_25, belonging to ANME-2c, also encode S-layer–associated multi-heme c-type cytochromes, implying a role of ANME-2c archaea with an S-layer protein in conducting electron derived from reverse methanogenesis shuttling from the archaeal membrane to the outside of the cell^46^. S11_12_8 and S11_6_25 affiliated with the family ANME-2c, also encode outer membrane cytochrome Z (*omcZ*) gene (**Table S9**) which plays an important role in Fe(III) reduction^47^. Furthermore, S11_12_8 not only actively expressed at zone C with the maximum of MAG’s abundance and TPM values with *mcrB* gene, but also had the significantly positive relation with the CH_4_ (ρ=0.790, *p*<0.01), Fe^2+^ (ρ=0.734, *p*<0.01), Mn^2+^ (ρ=0.601, *p*<0.05; **Figure 3b**).

Gene encoding decaheme c-type cytochrome (*mtrC*), was present in S11_6_22 and Co_S11_566 affiliated with ETH-SRB1 from the order *Desulfobacterales* (**Table S5**). The two MAGs (S11_6_22 and Co_S11_566) have the higher abundance in the methanic zone (Mean: 0.20% and 0.12%) than the SMTZ (Mean: 0.18% and 0.07%; **Table S3**). Besides, spearman’s correlation results (**Figure 3b**) show that Co_S11_566 closely related with concentrations of Fe^2+^ (ρ=0. 699) and Mn^2+^ (ρ=0. 650). Thus, ETH-SRB1 probably act as the role of metal reducing bacteria in the methanic zone. We also found the presence of hypothetical proteins attributed to porins, cytochrome *c* binding motif sites (CxxCH) and Geobacter-related gene markers (*omc*) for iron reduction in Co_S11_933 (**Table S9**), belonging to Zixibacteria, which was reported with pathways of either oxidation or reduction of ferric/ferrous iron and arsenate/arenite and nitrate/nitrite^48^. Co_S11_933 also displayed a higher abundance in the methanic zone (0.03∼0.06%) than in other zones (**Table S3**).

### Contribution of Metal-AOM to methane consumption

Geochemical observations and microbiological analyses support that Fe and Mn oxides reduction is coupled to methane oxidation in the methanic zone. We then used numerical modeling to predict their contributions to methane consumption. Constrained by the measured porewater data and Fe leaching experiments (**Figure 1; Tables S1, S6 and S7**), the results of the reaction-transport modeling predict the model-derived rates for Fe-AOM of up ∼0.02 μmol CH_4_ cm^−3^ d^−1^ in the methanic zone (**Figure 4 and Table 1**). These rates are more than 20 times as big as the estimated potential Fe-AOM rates from in situ marine methanic sediments with much higher Fe^2+^ concentration (180∼800 μM)^20-23^ (**Table 1**), but much lower than those derived from stimulated microbial communities in laboratory incubation studies5, 24, 25, 49.

**Figure 4.**
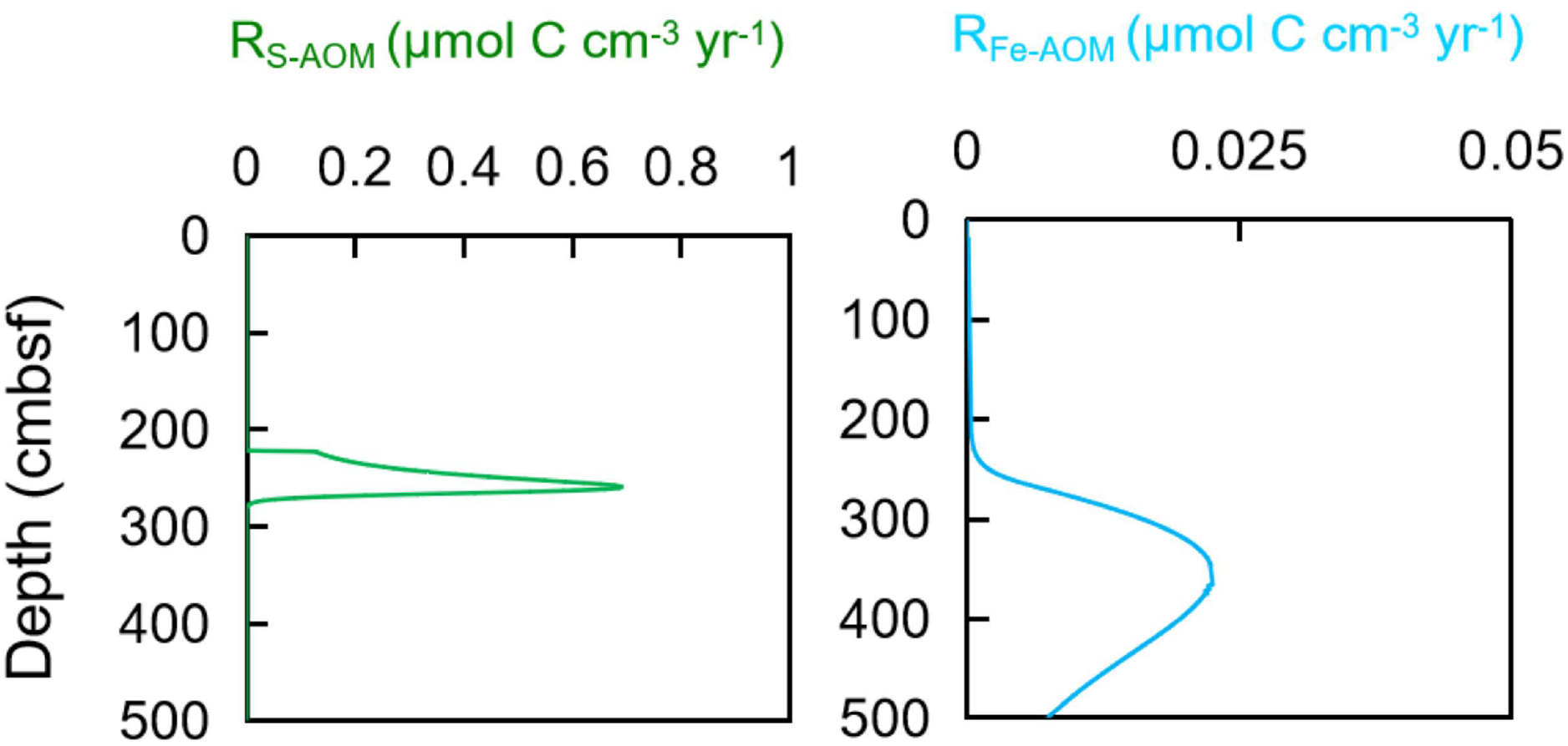
Modeled reaction rate profiles of S-AOM (green), and Fe-AOM (blue).

As Mn speciation data were not available, we used the diffusive Mn^2+^ flux calculated based on the quasi-linear concentration gradient at the depth interval of ∼250 cm and 350 cm to represent the rate of Mn-AOM. Based on the porewater profiles, our estimated diffusive flux for Mn^2+^ is 1.276 μmol cm^-2^ yr^-1^. Thus, taking into account that only one molecule of CH_4_ is needed to reduce four molecule of MnO_2_, CH_4_ removal by Mn-AOM is estimated to 0.319 μmol CH_4_ cm^-2^ yr^-1 19^. This is a minimum estimate as the potential Mn^2+^ consumption by authigenic minerals is not taken into account. Depth-integrated rates of Fe-AOM and Mn-AOM are both 0.3 μmol cm^-2^ yr^-1^ in the methanic zone, which are considerably lower than S-AOM rate (∼20 μmol cm^-2^ yr^-1^) and account for ∼1.5% of total CH_4_ removal by microbial metabolism, respectively (**Table 1**). The high S-AOM rate is caused by methane bubble dissolution while its upward-ascending and enhanced sulfate supply from seawater due to bubble irrigation. The estimated depth-integrated rate of Fe-AOM in the Haima seep falls within the range of those reported in different environments globally^20-23^. These data from Haima cold seep provide the first in situ evidence for quantitatively significant manganese-dependent AOM in marine sediments. Given an apparent elevated sedimentary manganese content in the methanic zone (from 0.04% to 0.06%) and high concentration of dissolved Mn^2+^ (up to 2289 μM), the contribution for Mn-AOM consumed by authigenic minerals could have been underestimated.

## Conclusions

Methane oxidation occurs in methanic zones driven by sedimentary microbial communities is an important mechanism that controls natural emissions of methane from the gas hydrate–bearing area. It happens mainly due to the presence of alternative electron acceptors other than sulfate to react with methane. Abundant Fe/Mn-(oxyhydr)oxides preserved in the SCS shelf sediments might be migrated into the study region due to the rapid increase of anomalous subsidence towards the deep water areas in the Qiongdongnan basins (**Figure 5**). Therefore, high amounts of buried reactive Fe(III)/Mn(IV) minerals seem to be an important available electron acceptors for AOM in the methanic zone of the Haima methane seep, accompanying by the generation of highly alkaline, extremely δ^13^C_DIC_-depleted and Fe(II)/Mn(II)-enriched pore waters, abundant Fe–Mn carbonates, along with authigenic magnetite by microbial iron/manganese reduction. In methanic sediments, abundant active ANME groups (ANME-1 and ANME-2c), and potential dissimilatory iron reducers (e.g. ETH-SRB1) are potentially involved in Metal-AOM *in situ*. Mechanistically, the apparent ability of ANME-2c to oxidize methane via the release of single electrons in this study should also be able to respire solid electron acceptors directly via extracellular metal reduction, which would confirm the presence of previously reported methane oxidation coupled to insoluble Fe(III)/Mn(IV) reduction. It is estimated that Metal-AOM at least contributes 3% CH_4_ removal from methanic sediments to the seep. Overall, Metal-AOM could significantly impact the biogeochemical cycles in consuming CH_4_ in modern marine sediments.

**Figure 5.**
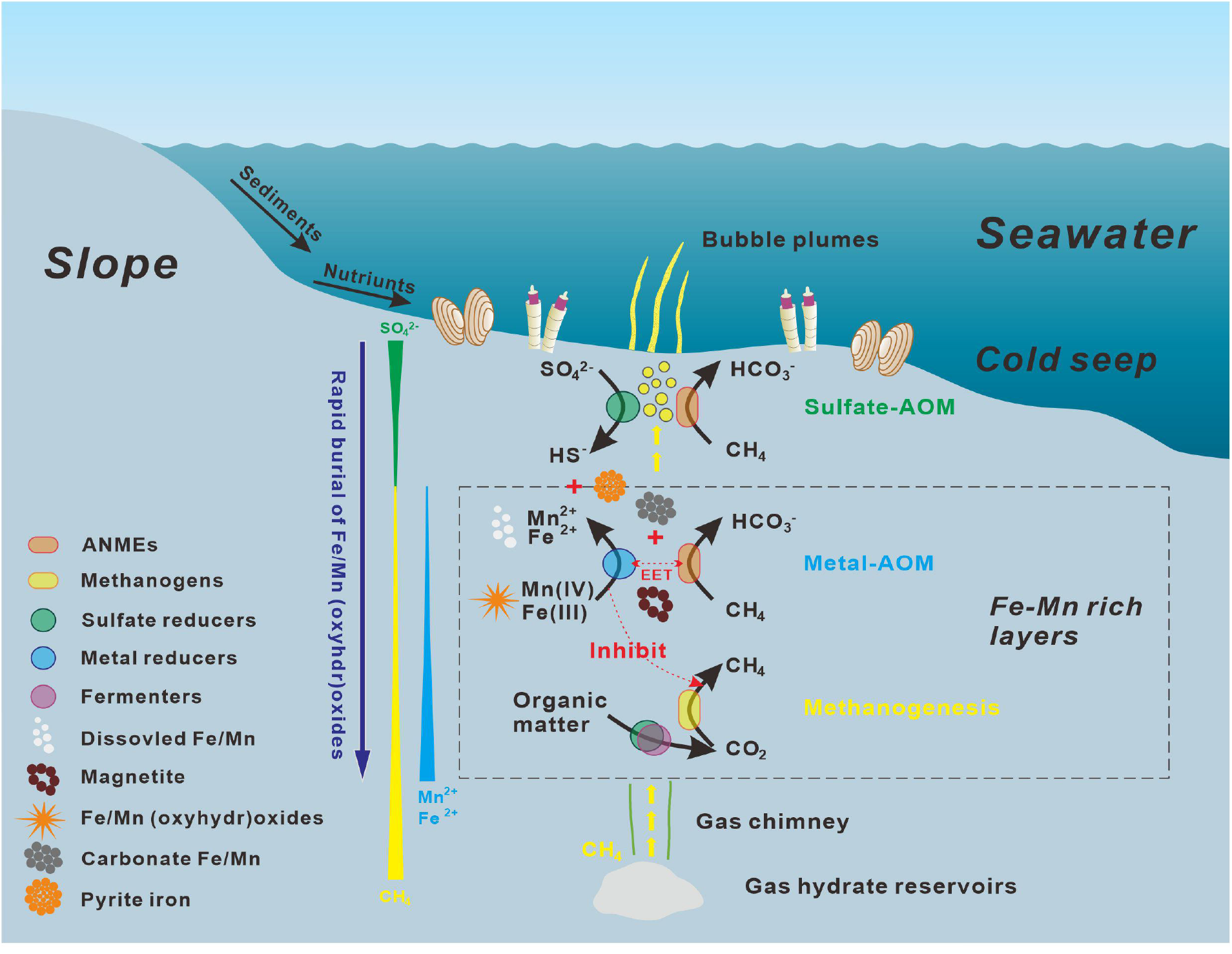
Model for AOM and methanogenesis in cold-seep marine sediments.

## Materials and methods

### Sampling and geochemical analyses

Sediment core HM-S11 with the length of 430 cm was obtained by a gravity piston sampler at the ROV1 site of Haima cold seeps^28^ (**Figure S1**) during the Haiyangdizhi10 cruise in June 2019 by the Guangzhou Marine Geological Survey, China Geological Survey.

The pore water was extracted from each sediment segment with the interval of 40 cm except the top 60 cm (20-cm interval) on board at room temperature by a vacuum apparatus. The concentrations of Fe^2+^ and Mn^2+^ in pore water were immediately on board determined by UV/Vis spectrophotometer (Beijing Purkinje, China) using the 1,10-phenanthroline colorimetric method and potassium periodate oxidation spectrophotometry, respectively. Sulfate concentrations were measured by an ICS-1100 ion chromatography (Thermofisher, USA) with an analytical error of ±1%. The δ^13^C_DIC_ values in pore water were analyzed by a multiflow-isotope ratio mass spectrometer (Thermofisher, Delta V Advantage, USA), with an analytical error of ±0.2‰. The concentrations of PO_4_^3−^ and NH_4_^+^ were photometrically measured on board using a UV-Vis spectrophotometer (Hitachi U5100, Hitachi Limited, Tokyo, Japan) with an analytical error of ±3.0%. Porosity was determined from the weight loss before and after freeze-drying of the wet sediments^37^ using a cutting ring with the volume of 5 ml on board, assuming a density of the porewater of 1.0 g cm^-3^.

The C_1_∼C_3_ concentrations of the headspace gas samples were determined using an Agilent 6850 gas chromatograph (Thermo, USA) with a Porapark Q column. The detection limit for all gases is 10□ppm, and the quantification limit is 30□ppm. The δ^13^C values of the methane were measured using gas chromatography-isotope ratio-mass spectrometry (GC-IR-MS; Thermo, USA), and are reported relative to the Vienna Peedee Belemnite standard (V-PDB), with an analytical error of ±0.5‰.

The major element composition of sediments was determined by an iCAP 7200 Inductively Coupled Plasma (ICP) Optical Emission Spectrometry (OES) (Thermo, USA). The content of different iron phase mineral components was determined by Infinite M200Pro multifunction enzyme marker (TECAN, USA) with a sequential extraction method^50^, and measured at the light absorption wavelength of 510 nm. The accuracy and repeatability of the absorption wavelength of the instrument are < ±0.5 nm (λ > 315 nm).

## DNA and RNA extraction

Total genomic DNA and RNA of each sample was extracted using Soil DNA Kit and Soil RNA Mini Kit (Omega Bio-Tek Inc., Norcross, GA) according to the manufacturer’s instructions, respectively. DNA concentration and purity were measured by TBS-380 (Turner Biosystems, CA, USA) and Nanodrop ND-2000 (Thermo Fisher Scientific, Waltham, USA), respectively. DNA extract quality, RNA degradation and contamination were monitored on 1% agarose gels. RNA quantity was measured using Qubit 2.0 (Thermo Fisher Scientific, MA, USA) and Nanodrop One (Thermo Fisher Scientific, MA, USA) at the same time. RNA integrity was accurately detected using the Agilent 2100 system (Agilent Technologies, Waldbron, Germany).

### Amplicon analysis of 16S rRNA genes and transcripts

The DNA and RNA for each sample were amplified in triplicate using primers 338F/806R for bacteria and Arch344F/Arch915R for archaea. Their PCR products were pooled and purified. The same amount of the PCR product from each sample was mixed to construct a sequencing library. High-throughput sequencing was carried out on the Illumina MiSeq sequencing platform using PE300 chemical at Majorbio Bio-Pharm Technology Co. Ltd. (Shanghai, China).

After demultiplexing, the resulting sequences were merged with FLASH (v1.2.11)^51^ and quality filtered with fastp (v0.19.6)^52^. Then the high-quality sequences were denoised using DADA2^53^ plugin in the Qiime2 (v2020.2)^54^ pipeline with recommended parameters, which obtains single nucleotide resolution based on error profiles within samples. DADA2 denoised sequences are usually called amplicon sequence variants (ASVs). To minimize the effects of sequencing depth on alpha and beta diversity measure, the number of sequences from each sample was rarefied to 4000, which still yielded an average Good’s coverage of 97.90%. Taxonomic assignment of ASVs was performed using the Blast consensus taxonomy classifier implemented in Qiime2 and the SILVA 16S rRNA database (v138).

### Metagenomic sequencing

DNA extract was fragmented to an average size of about 400 bp using Covaris M220 (Gene Company Limited, China) for paired-end library construction. Paired-end library was constructed using NEXTFLEX^®^ Rapid DNA-Seq (Bioo Scientific, Austin, TX, USA). Adapters containing the full complement of sequencing primer hybridization sites were ligated to the blunt-end of fragments. Paired-end sequencing was performed on Illumina NovaSeq (Illumina Inc., San Diego, CA, USA) at Majorbio Bio-Pharm Technology Co., Ltd. (Shanghai, China) using NovaSeq Reagent Kits according to the manufacturer’s instructions (www.illumina.com).

### Assembly and binning of metagenomes

Raw reads derived from the 13 metagenome libraries were quality-controlled by clipping off primers and adapters then filtering out artefacts and low-quality reads using Read_QC module within the metaWRAP pipeline v1.3.2 ^55^. Filtered reads were individually assembled using SPAdes v3.13.0 (metaSPAdes mode, default parameters, for samples S11_2-3, S11_8-10, S11_12-13)^56^ or MEGAHIT v1.1.3 (default parameters, for samples S11_1, S11_4-7, S11_11)^57^. Additionally, all samples were co-assembled using MEGAHIT v1.2.9 (--kmin_1pass --k-min 31). Each assembly was binned using the binning module within the metaWRAP pipeline v1.3.2 (--metabat2 --maxbin2 --concoct for individual assembly; --metabat2 for co-assembly). For each assembly except S11_10, the three bin sets (one for co-assembly) were then consolidated into a final bin set with the bin_refinement module of metaWRAP pipeline v1.3.2 (-c 50 -x 10). For S11_10, bin sets were consolidated into a final bin set with DAS Tool v1.1.2 (default parameters)^58^. Finally, 638 bins were obtained from the 14 assemblies. They were then combined and dereplicated using dRep v 2.6.2 (-comp 50 -con 10 -sa 0.95 --S_algorithm fastANI) at 95% average nucleotide identity clustering (species level)^59^. After dereplication, a total of 351 dereplicated MAGs were obtained. Each bin was taxonomically assigned according to the Genome Taxonomy Database (GTDB) version r207 using GTDB-tk v2.0.0 ^60^.

MAGs were annotated with FeGenie v2.0 and METABOLIC v4.0 ^61, 62^. CoverM v0.5.0 “genome” (https://github.com/wwood/CoverM) was used to obtain relative abundance of each genome (parameters: –min-read-percent-identity 0.95 –min-read-aligned-percent 0.75 –trim-min 0.10 –trim-max 0.90). For *mcr*A phylogenetic analyses, amino acid sequences were aligned with MUSCLE algorithm^63^ (-maxiters 16) and the maximum-likelihood phylogenetic tree was constructed in MEGA X^64^.

### Metatranscriptomic sequencing

Whole mRNAseq libraries were generated by Guangdong Magigene Biotechnology Co.,Ltd. (Guangzhou, China) using NEB Next^®^ Ultra™ Nondirectional RNA Library Prep Kit for Illumina^®^ (New England Biolabs, MA, USA) following manufacturer’s recommendations. Briefly, the bacterial and archaeal 16S rRNA transcripts in total RNA samples were reduced by Ribo-zero rRNA Removal Kit. Fragmentation was carried out using NEB Next First Strand Synthesis Reaction Buffer. The first strand cDNA was synthesized using random hexamer primer and M-MuLV Reverse Transcriptase (RNase H), and the second strand cDNA synthesis was subsequently performed using DNA Polymerase I and RNase H. Remaining overhangs were converted into blunt ends via exonuclease/polymerase activities. After adenylation of 3’ ends of DNA fragments, NEB Next Adaptor with hairpin loop structure were ligated to prepare for hybridization. In order to select cDNA fragments of preferentially 150∼200 bp in length, the fragments were selected with AMPure XP beads (Beckman Coulter, Beverly, USA). Then PCR was performed with Phusion High-Fidelity DNA polymerase, Universal PCR primers and Index (X) Primer. At last, PCR products were purified with AMPure XP beads and library insert size was assessed on the Agilent 2100 system (Agilent Technologies, Waldbron, Germany). The clustering of the index-coded samples was performed on a cBot Cluster Generation System. After cluster generation, the library was sequenced on an Illumina Hiseq Xten platform and 150 bp paired-end reads were generated.

### Metatranscriptomic analysis

Raw metatranscriptomic reads were quality filtered using Read_QC module within the metaWRAP pipeline v1.3.2 ^55^. Paired-end reads were merged using PEAR v0.9.8 ^65^. The reads corresponding to ribosomal RNAs were removed using SortMeRNA v4.2.0 ^66^. Subsequently, mRNA reads were mapped to the predicted protein-coding genes from MAGs using Salmon v1.5.0 in mapping-based mode (parameters: - validateMappings -meta)^67^. The expression level for each gene was normalized to transcript per million (TPM).

### Numerical modeling

A one-dimensional, reaction-transport model^22, 68^ was applied to simulate two solid phases (reactive Fe oxides and Fe carbonate) and six dissolved species (SO_4_^2-^, CH_4_, DIC, Ca^2+^, Mg^2+^, and Fe^2+^). The reactions and their kinetic rate expressions considered in the model are listed in **Table S1**. Net reaction terms for all species and model parameters are listed in **Tables S11 and S12**, respectively. Detailed model construction can be found in Supplementary Materials.

### Flux calculations

Diffusive fluxes *J* (μmol cm^-2^ yr^-1^) of dissolved Mn^2+^ in the methanic zone were calculated using its pore water concentration gradients according to Fick’s first law of diffusion. The algorithms are detailly described in Supplementary Materials.

### Statistical analysis

The Spearman’s linear correlation among the geochemical parameters and microbial abundances of the subsamples was analyzed using the R package Vegan^69^ via RStudio (Ver. 1.3.1093).

### Data availability

All metagenomes and metatranscriptomes are available at the Sequence Read Archive under BioProject accession number PRJNA738468. Amplicon data were deposited in Figshare (https://doi.org/10.6084/m9.figshare.21696296.v1).

## Acknowledgements

This work was supported by the Guangdong Basic and Applied Basic Research Foundation (No. 2019B030302004, 20201910240000691), and the National Natural Science Foundation of China (No. 41906076, 41806074, 41730528), the Marine Geological Survey Program of China Geological Survey (DD20221706). The authors express their sincere gratitude to the crews and participates of the Haiyangdizhi10 for assistance in collecting, processing, and shipping the samples.

## Author contributions

QL and XD designed the study. XX, TZ, and XW performed porewater and sediment sampling, geochemical analyses. ML performed geochemical modeling estimates. XD and XX processed metagenome, metatranscriptome data analyses. JT and ZC provided the picture of seafloor observations. XW and JZ interpreted geophysical data. XX and TZ conducted amplicon sequencing and microbial diversity analyses. XX, XD, and ML wrote the manuscript. QL, CZ, and XY modified the manuscript. All authors reviewed the results and approved the submitted manuscript.

