## Supplementary Materials for "Metal-driven anaerobic oxidation of methane as an important methane sink in methanic cold seep sediments"

### Numerical modeling

Major reactions considered in the model are SO_4_^2-^-AOM, Fe-AOM, and authigenic carbonate precipitation. Organic matter degradation which is thought to play a minor in seep sediments is not considered in the model. This assumption is plausible given very low pore-water NH_4_^+^ and PO_4_^3-^ concentrations. Two partial differential equations were used to resolve the depth-concentration profiles of solid and dissolved species, respectively ^1, 2^:

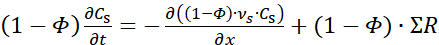
 (S1)

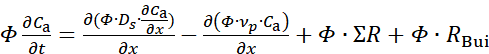
 (S2)

where *x* (cm) is depth in the sediments, *t* (yr) is time,
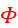
 (dimensionless) is porosity, *C*_s_ ( dry wt. %) is the solid content, *C*_a_ (mmol cm^-3^) is the concentration of dissolved species, *D_s_* (cm^2^ yr^-1^) is the molecular diffusion coefficient corrected for tortuosity,
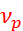
 (cm yr^-1^) is the burial velocity of porewater,
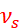
 (cm yr^-1^) is the burial velocity of solids, Σ*R* denotes the sum of the rates of biogeochemical reactions considered in the model, and *R*_Bui_ is the mixing rate of bottom water and porewater by bubble irrigation.

Sediment porosity decreases with depth assuming steady-state compaction:

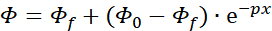
 (S3)

where
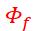
 and
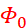
 (both dimensionless) denote the porosity below the depth of compaction and at sediment surface, respectively, and *p* (cm^-1^) is the porosity attenuation coefficient.

In the absence of externally-imposed fluid advection at the seafloor, the velocity of porewater and solids is directed downward under steady-state compaction relative to the seafloor:

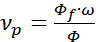
 (S4)

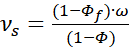
 (S5)

where
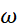
 (cm yr^-1^) is the sedimentation rate taken from Hu et al. (2019)^3^. Depth-dependent molecular diffusion coefficients of dissolved species were calculated after Boudreau (1997)^4^ and Oelkers and Helgeson (1991)^5^ and corrected for tortuosity:

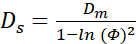
 (S6)

where *D_m_* is the molecular diffusion coefficient in free seawater at the in-situ temperature, salinity and pressure. The diffusive transport of DIC was simulated using the diffusion coefficient of bicarbonate (HCO_3_^-^) since this is the dominant anion.

Rising gas bubbles facilitate the exchange of porewater and bottom water as they move through tube structures in soft sediments^6^. The induced porewater mixing process was described as a non-local transport mechanism whose rate for each species is proportional to the difference between solute concentrations at the sediment surface *C_0_* (mmol cm^-3^) and at depth below the sediment surface *C_x_* (mmol cm^-3^) (*R*_Bui_, Table S1). Bubble irrigation is described by parameters *α_1_* (yr^-1^) and *α_2_* (cm) that define (respectively) the irrigation intensity and its attenuation below the irrigation depth *L*_irr_ (cm) ^7^.

The rate of gas dissolution, *R*_diss_ (mmol cm^-3^ yr^-1^), was described using a pseudo first-order kinetic expression of the departure from the local methane gas solubility concentration, *L*_MB_ (mmol cm^-3^), where *k*_MB_ (yr^-1^) is the kinetic constant for gas bubble dissolution (**Table S12**). Methane only dissolves if the porewater is undersaturated with respect to *L*_MB_:

CH_4_(g) → CH_4_(aq)	for CH_4_ ≤ *L*_MB_	(S7)

*L*_MB_ was calculated for the in-situ salinity, temperature and pressure using the algorithm in **Duan et al. (1992) ^8^**. *k*_MB_ was constrained using the dissolved sulfate and DIC data.

Methane consumed by both SO_4_^2-^-AOM (CH_4_ + SO_4_^2-^ → HCO_3_^-^ + HS^-^ + H_2_O) and Fe-AOM (CH_4_ + 8Fe(OH)_3_ + 15H^+^ → HCO_3_^-^ + 8Fe^2+^ + 21H_2_O) is calculated using bimolecular kinetics:

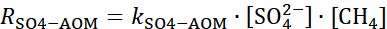
 (S8)

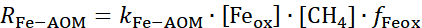
 (S9)

Where k_SO4-AOM_ (yr^-1^) and k_Fe-AOM_ (yr^-1^) represent rate constants for SO_4_^2-^-AOM and Fe-AOM, respectively. f_Feox_ is the unit conversion term between reactive Fe oxides (dry wt.%) and Fe^2+^ (mmol cm^-3^).

The rates of Ca^2+^, Mg^2+^, and Fe^2+^ consumed by carbonate precipitation were estimated using an inverse modeling procedure, through which the rates were obtained without specifying the kinetic rate law for authigenic carbonate precipitation. For Ca^2+^ and Mg^2+^, empirical functions were first fitted to the measured Ca^2+^ and Mg^2+^ concentrations to obtain best-fit concentration profiles of observed Ca^2+^ (Ca_OBS_) and Mg^2+^ (Ca_OBS_). The net reaction rates of Ca^2+^ and Mg^2+^ (mmol dm^-3^ yr^-1^) were set proportional to the difference between simulated and observed concentrations:

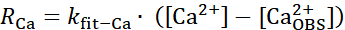
 (S10)

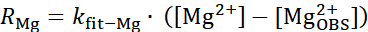
 (S11)

The net Fe^2+^ release rate was calculated using the same approach:

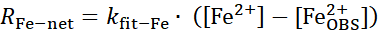
 (S12)

where *k*_fit-Ca_ (yr^-1^), *k*_fit-Mg_ (yr^-1^) *k*_fit-Fe_ (yr^-1^) is the kinetic rate constants with a high value to ensure that the modeled concentrations were maintained close to the measured values.

As Fe^2+^ is produced by Fe-AOM and consumed by Fe carbonate precipitation, the rate of Fe consumption by authigenic carbonate precipitation (R_Fe-Carb_, mmol dm^-3^ yr^-1^) was given by:

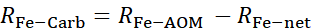
 (S13)

The length of the simulated domain was set to 500 cm. All the parameters applied in the model were listed in Table S12. Upper boundary conditions for all dissolved species were imposed as fixed concentrations (Dirichlet boundary) using measured values in the uppermost sediment layer. Upper boundary conditions for solid phases (Fe_ox_ and Fe_CaCO3_) were prescribed as fluxes (Robin-type boundary). A zero-concentration gradient (Neumann-type boundary) was imposed at the lower boundary for all species. The continuous differential equations in Eqs. (S1) and (S2) were solved using finite differences and the method-of-lines over an uneven grid with a high spatial resolution at the surface and lower resolution toward the bottom. The model was solved using the NDSolve object of MATHEMATICA V. 9.0. Mass conservation of >99 % was reached for all simulations.

### Flux calculations

Assuming that all of the Mn-reduction below the SMTZ is due to Mn-AOM and limited removal of dissolved Mn^2+^ below the SMTZ, diffusive fluxes *J* (μmol cm^-2^ yr^-1^) of dissolved Mn^2+^ in the methanic zone were calculated as follows (Eq. S14):

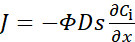
 (S14)

where
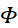
 is the measured porosity (**Table S1**), *D_s_* (cm^2^ yr^-1^) is the molecular diffusion coefficient corrected for tortuosity,
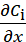
 (μmol cm^-4^) is the solute concentration gradient, which was estimated based on the linear portions of concentration profiles within the investigated depth intervals. Sediment diffusion coefficients were calculated as follows:

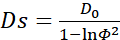
 (S15)

where *D_0_* (diffusive flux of Mn^2+^ in seawater, cm^2^ s^-1^) was corrected for in-situ temperature and salinity (5 ºC, 3.75×10^-11^ m^2^ s^-1^)^4^.

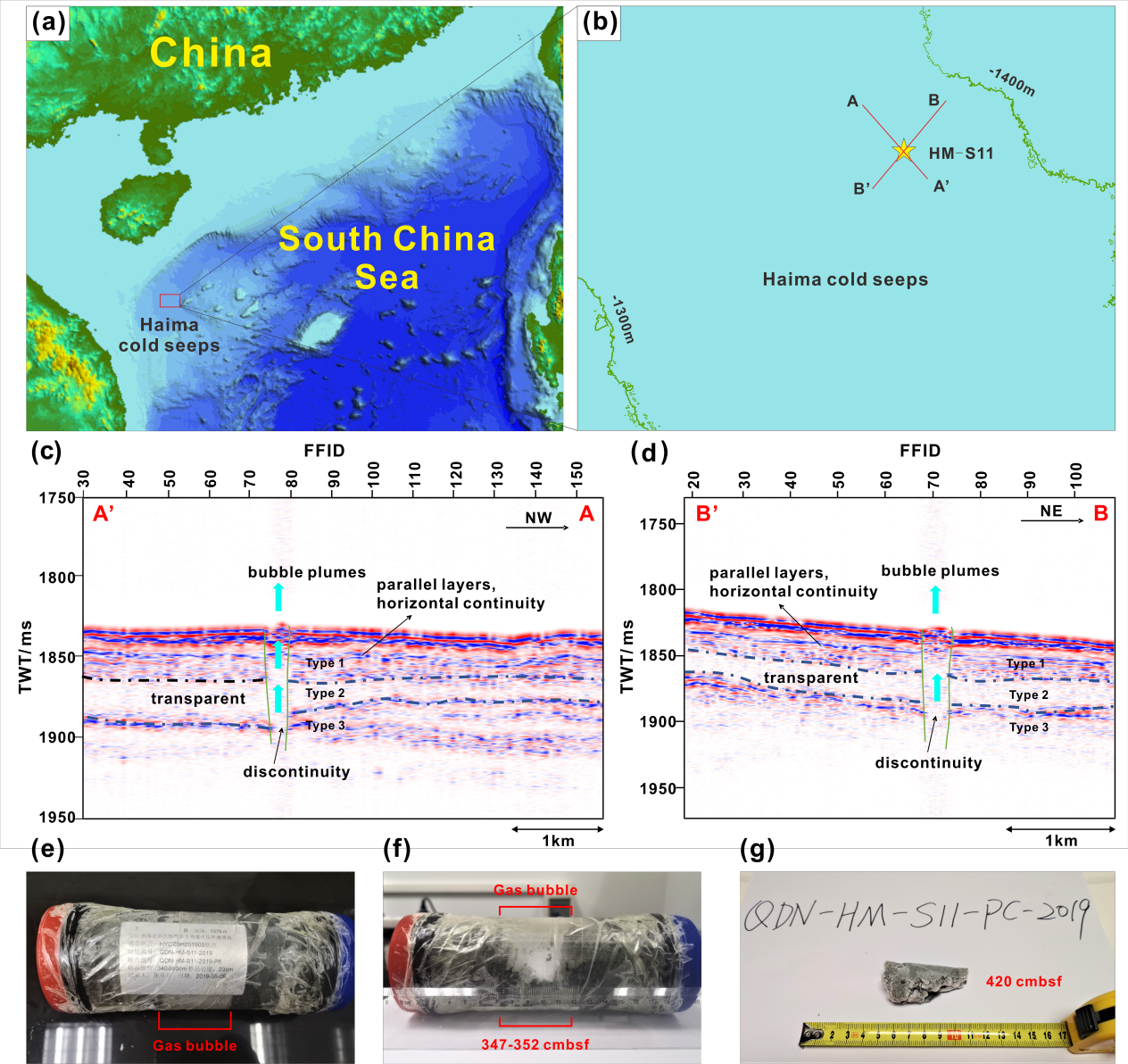

**Figure S1.** **Maps showing the location of the sediment core HM-S11.** (a) Bathymetry map of the study area. (b) The direction of sesimic lines; the intersection of two lines is over-leakage point; the Line A'A is from southeast (SE) to northwest (NW); the Line B'B is from southwest (SW) to northeast (NE). Seismic profiles in (c) A'A and (d) B'B indicate the gas and fluid migration pathways from deep sediments to the seabed (cyan arrows). The horizontal axis "FFID" means Field File ID; The vertical axis "TWT" is two-way travel time, where "ms" stands for milliseconds. (e) and (f) show gas bubbles in the depth of 340-360 cmbsf of the piston core. (g) Massive gas hydrates were found at 420 cmbsf in the bottom of the piston core.

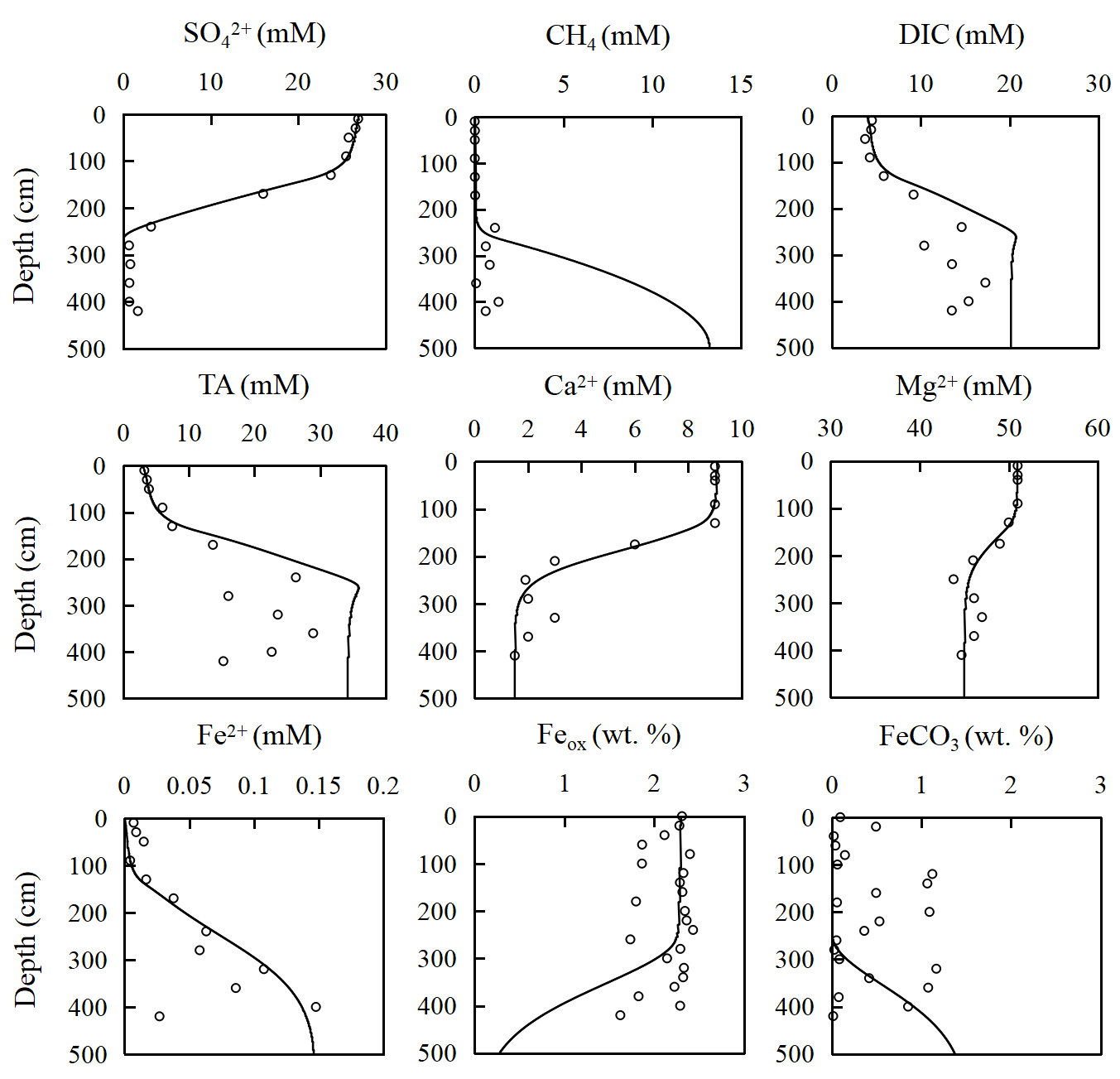

**Figure S2** Measured data (dots) and simulated results from the steady-state model (curves) of core HM-S11. Down-depth concentration of sulfate (SO_4_^2-^), methane (CH_4_), dissolve inorganic carbon (DIC), total alkalinity (TA), calcium (Ca^2+^), magnesium (Mg^2+^), dissolved iron (Fe^2+^), iron (oxyhydr)oxides (Fe_ox_), carbonate-associated Fe (FeCO_3_), are shown.
